# Over-Confluence of expanded bone marrow mesenchymal stem cells ameliorates their chondrogenic capacity in 3D cartilage tissue engineering

**DOI:** 10.1101/2020.01.08.897645

**Authors:** Damien Tucker, Karen Still, Ashley Blom, Anthony P. Hollander, Wael Kafienah

## Abstract

Cartilage tissue engineering using bone marrow-derived mesenchymal stem cells (BM-MSCs) is a growing technology for the repair of joint defects. Culturing BM-MSCs to over confluence has historically been avoided due to perceived risk to cell viability, growth inhibition and differentiation potential. Here we show that a simple change in culture practice, based on mimicking the condensation phase during embryonic cartilage development, results in biochemically and histologically superior cartilage tissue engineered constructs. Whole transcriptome analysis of the condensing cells revealed a phenotype associated with early commitment to chondrogenic precursors. This simple adjustment to the common stem cell culture technique would impact the quality of all cartilage tissue engineering modalities utilising these cells.

## INTRODUCTION

Cartilage tissue engineering strategies have sought a cell population capable of enhanced chondrogenesis that could be reliably used in regenerative medicine strategies (1). Mesenchymal stem cells (MSCs) taken from the bone marrow are a very commonly used cell type for such strategies. These cells however suffer from heterogeneity; a lack of consistent culture methods leads to variable outcomes in cartilage repair approaches (2, 3). Many strategies have been explored to improve the consistency of MSC-based cartilage regeneration including clonal expansion based on cell surface markers (e.g. STRO-1 (4), CD271 (5) and ROR2 (6)). Whilst such approaches have showed promise they remain inefficient and expensive.

Scaling up tissue engineering strategies for regenerative medicine applications will require a large number of cells invariably taken through multiple doublings. Historically, MSC cell culture methodologies have avoided growing MSCs to over confluence since it was hypothesised it can lead to decreased cellular viability and differentiation potential (7, 8). Increased confluence or cell density is found during embryonic chondrogenesis when previously dispersed mesenchymal cells condense to differentiate into chondrocytes – the earliest sign of skeletal development through endochondral ossification (9, 10). *In vitro* culture of BM-MSCs to over confluence can recapitulate cell condensation behaviour *in vivo* prior to chondrogenesis.

Here we demonstrate for the first time that BM-MSCs cultured to over confluence not only maintain their cellular viability but are able to produce three-dimensional cartilage constructs of superior quality than cells cultured by traditional methods. This simple methodological change in culture will facilitate enhanced *in vitro* cartilage production using BM-MSCs for a wide range of cartilage cell-based therapies.

## RESULTS & DISCUSSION

### BASIC CHARACTERISATION OF OVER CONFLUENT MSC’S

The growth kinetics and multipotency of over confluent BM-MSCs was assessed at multiple time points and over a number of passages. This was performed in order to ascertain whether BM-MSCs grown to over confluence maintain their ‘stemness’. BM-MSCs from 5 individual donors were plated at a known cell density (2000 cells/cm^2^) in multiple culture flasks and harvested at set time points to determine the cell density, cellular viability and population doubling time. Cell density analysis demonstrated a gradual increase in confluence over 24 days post plating in passage 2 cells (Fig. 1A). Visual confirmation of confluence was attained in all populations by 10 days (Supplementary Fig 1).

**Figure 1:**
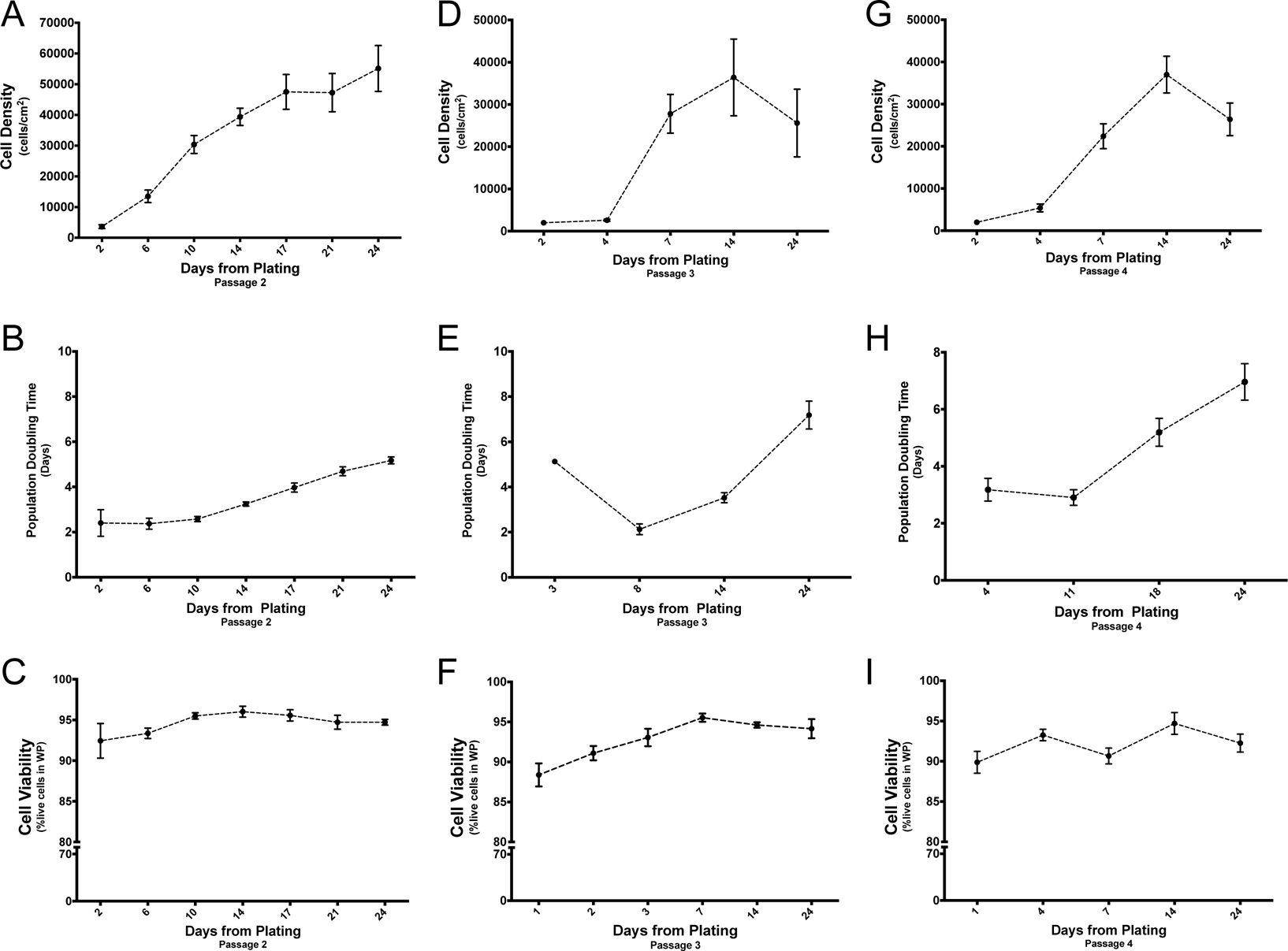
Basic characterisation of over confluent MSC’s. BM-MSC cell lines from 5 individual donors from passage 2 were plated at a cell density of 2000 cells/cm^2^ and permitted to proliferate in expansion media supplemented with rh-FGF. At defined time points the cells were harvested, counted and analysed using flow cytometry to determine the cell density at harvest (A, D & G), the population doubling time (B, E & H) and the cell viability expressed as a percentage of the WP at flow cytometry (C, F & I). The cells harvested after 24 days in culture were analysed and re-plated at a cell density of 2000 cells/cm^2^ and permitted to proliferate as passage 3 cells and analysed as before (D, E & F); this procedure was repeated for passage 4 cells (G, H & I).

The population doubling time – a measurement of cellular proliferation - increased as cell density increases (Fig 1B). Flow cytometry analysis of harvested cells - utilising propidium iodide (PI) as a viability marker - revealed viability in excess of 90% at all time points (Fig. 1C). Further evidence of maintained cellular viability in over confluent cells was demonstrated by the continued proliferation of over confluent cells following re-plating. Over confluent cells re-plated at a low cell density demonstrated an initial lag in proliferation (Fig. 1A, D, G). Following 4 days in culture, the population doubling time decreased, correlating with an increase in cell density and confluence (Fig 1D&E). Cell viability was less than the previous passage for the first few days returning to pre-plating levels after 3 days in culture. Cellular viability remained in excess of 85% in all populations at all time points in passage 3 (Fig 1E).

Passage 3 cells cultured for 24 days were again re-plated to assess proliferation and viability. These cells were able to proliferate at a similar rate to cells from previous passages and maintain viability in excess of 90% at most time points (Fig 1H&I). Increase in cell density followed a similar pattern to those cells in passage 3 with a gradual increase to 14 days in culture followed by a slight decrease in cell density at 24 days (Fig 1G). The growth curve of over confluent cells is linear and extended at passage 2 compared to replated over-confluent cells at later passages.

In order to assess the multipotency of over confluent BM-MSCs, 6 individual cell lines were permitted to reach over confluency and (after 24 days in culture) were stimulated to differentiate into adipogenic and osteogenic lines with the appropriate growth factors. The resulting monolayer cultures along with the corresponding undifferentiated controls were stained and photographed using bright field microscopy (Supplementary Fig 2). All cell lines underwent adipogenic and osteogenic differentiation confirming the maintenance of multipotency in over confluent cells. The adipogenic and osteogenic potential of these cell lines at low confluence (8 days in culture) was confirmed and appeared similar to over confluent cultures (Supplementary Fig 2).

**Figure 2:**
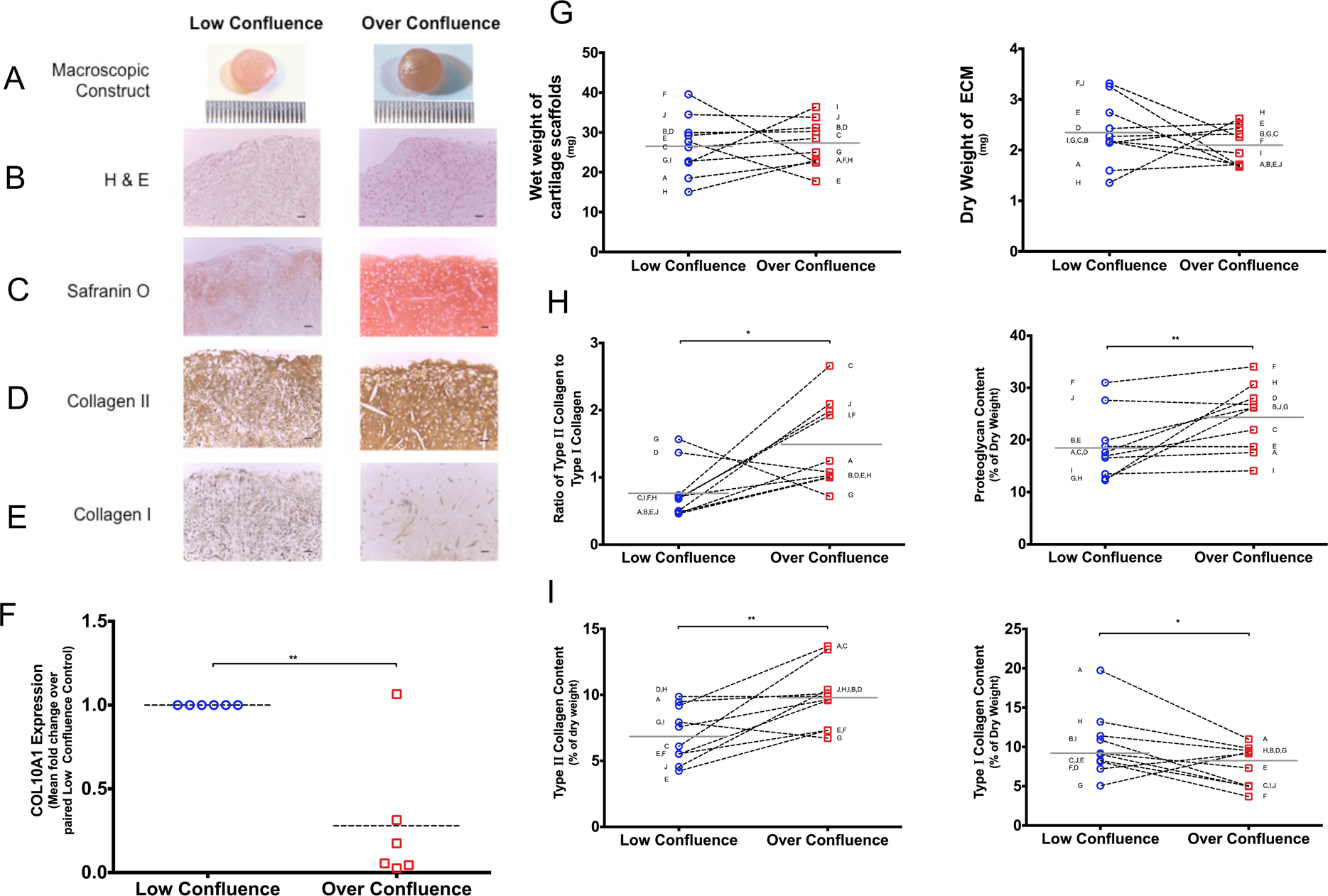
Over confluent BM-MSCs generate improved tissue-engineered cartilage. BM-MSCs from 10 individual cell lines were permitted to proliferate in expansion medium for 5 days (low confluent) and 24 days (over confluent) before being harvested and seeded onto PGA scaffolds coated with fibronectin and stimulated to undergo chondrogenesis in the presence of TGF-β3. Following 35 days in culture these scaffolds were harvested, stained for histology, digested for biochemical analysis and analysed for gene expression to determine the quality of the cartilage constructs produced. A – Representative macroscopic image of cartilage constructs. Scale bar represents 1mm. B – Histological image of representative sample demonstrating cellularity utilising H&E stain. C – Safranin O stain for proteoglycan content in representative histological image. D&E – Representative immunohistological stain demonstrating the type II collagen content (D) and the type I collagen content (E) present in the ECM. Scale bar corresponds to 100 μm. Original magnification 10x. F – The cartilage constructs were analysed for type X collagen gene expression using RT-qPCR with Taqman^®^ as the fluorescence reporter. Each sample was processed in triplicate with beta actin used as the control gene. Comparative quantification was achieved using the ΔΔCT_t_ method. G – Following harvest – at 35 days – the cartilage constructs were weighed to determine initially the wet weight of the cartilage constructs before the constructs were freeze dried to determine the dry weight of the ECM produced in each scaffold. Each point represents each individual cell line with the line depicting the mean for each culture condition. H&I – Duplicate three-dimensional tissue engineered cartilage scaffolds generated from paired BM-MSCs from low confluent and over confluent states were freeze dried, digested in trypsin and analysed for the presence of collagen type II, collagen type I and proteoglycan in the ECM. The collagen content of the ECM was determined using a reverse ELISA assay and reported as a ratio of type II collagen to type I collagen. A colourmetric assay was used to determine the proteoglycan content of the scaffolds and reported as a percentage of the dry weight of the scaffolds. Statistical analysis was performed using a paired t test. * = p<0.05, ** = p<0.01.

In summary, confluence is defined as a visual measurement of the fraction of growth area covered by cells while cell density provides a quantification of proliferation, albeit as a result of detachment or destructive assays (12). Cell culture methodology has historically avoided cellular over confluence as it is believed that over confluence negatively affects cell viability and differentiation (7, 8). This study demonstrated that cell viability in excess of 90% was maintained in each cell line cultured up to 24 days proving that cellular viability is not negatively affected by over confluence as was once assumed. Furthermore, trilineage potential was maintained in these cells demonstrating the ability of over confluent BM-MSCs to maintain their stem cell properties after confluence is reached. To our knowledge this is the first study to demonstrate that BM-MSCs cultured to over confluence maintain both cellular viability and trilineage potential.

### OVER CONFLUENT BM-MSCS GENERATE IMPROVED TISSUE-**ENGINEERED CARTILAGE**

We hypothesised that over confluent MSCs could recapitulate early embryonic events associated with active chondrogenesis and therefore an improved tissue engineered cartilage. To test this hypothesis, three-dimensional cartilage tissue engineering was performed using BM-MSCs from 10 individual cell lines harvested at low confluence (5 days) and over confluence (24 days) and stimulated to undergo chondrogenesis in the presence of TGF-β3. These constructs were then analysed to determine the quality of the cartilage produced using histology, biochemical assays and RT-qPCR gene expression quantification. All 10 cell lines produced constructs resembling cartilage at a macroscopic level; no macroscopic difference was observed between the paired constructs from low and over confluent cells (Fig 2A and Supplementary Fig 3). This observation was confirmed by the negligible difference in the wet weight of the cartilage scaffolds following harvest and the similar dry weights of the ECM in the lyophilised constructs (Supplementary Fig 3B). The constructs generated using over confluent cells produced significantly superior cartilage ECM. Histological analysis of the constructs revealed an ECM that stained extensively for proteoglycan and type II collagen and weakly for type I collagen in the over confluent constructs (Fig 2B, C, D & E). Extensive quantitative analysis of the engineered cartilage ECM was performed to determine the biochemical composition of the cartilage constructs utilising well-validated assays (14).

**Figure 3:**
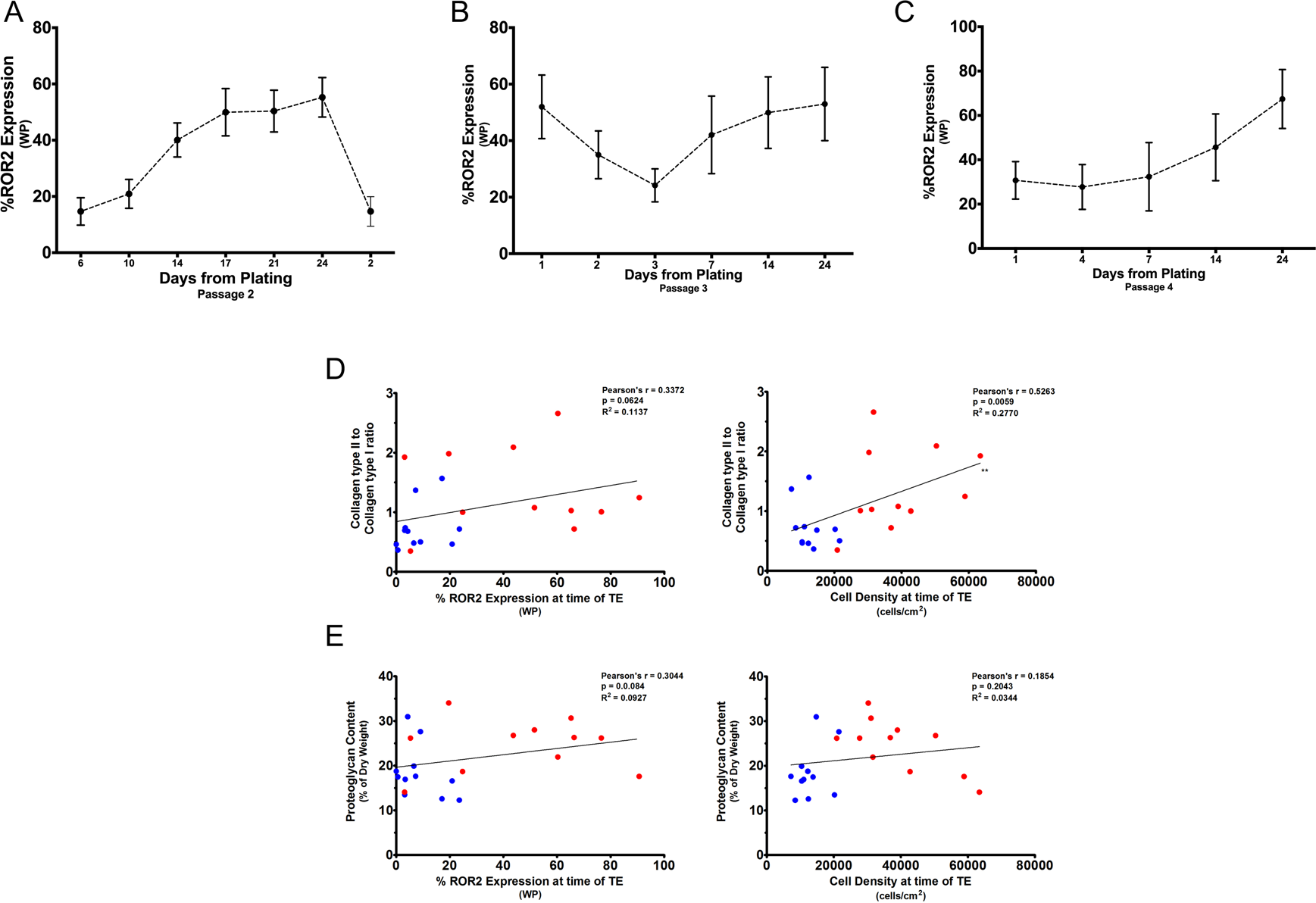
Over confluence leads to the enrichment of the highly chondrogenic ROR2-positive MSC population. The Wnt5a receptor ROR2 has been shown to be predictive of enhanced chondrogenesis in BM-MSCs. BM-MSCs from 5 individual cell lines were plated at a cell density of 2000 cells/cm^2^ and permitted to proliferate in expansion media before being harvested, counted and interrogated using flow cytometry to determine the percentage positivity of the marker ROR2 in the WP at defined time points until 24 days in culture. A – The WP ROR2 percentage is directly proportional to the number of days in culture increasing to 24 days from plating B – The cells harvested after 24 days were then re-plated at a cell density of 2000 cells/cm^2^ to determine the effect of re-plating on WP ROR2 positivity. C – The process was repeated for a further passage. Representative line graphs depicting the mean (and SEM) ROR2 expression in the whole population of cells at each time point. Prior to tissue engineering, low confluent (5 days in culture) and over confluent (24 days in culture) BM-MSCs from 10 paired individual cell lines were analysed using flow cytometry to determine the WP ROR2 positivity prior to tissue engineering. Biochemical analysis was performed on the generated cartilage constructs to quantify the quality of the ECM matrix produced. D – The ratio of type II collagen to type I collagen was plotted against the corresponding WP ROR2 percentage and cell density for that cell line line prior to the commencement of tissue engineering. E The proteoglycan content of the ECM produced by each construct was plotted against the corresponding WP ROR2 percentage and cell density for each cell line. Statistical analysis was performed using linear regression and Pearson’s correlation coefficient with an alpha value of 0.05 taken to be significant. ** = p<0.005.

Paired analysis comparing low confluent and over confluent constructs demonstrated a significantly increased type II collagen to type I collagen ratio (TII/TI ratio) and proteoglycan content in over confluent constructs (Fig 2H). A 2-fold increase in the TII/TI ratio was seen in the over confluent constructs while a 1.3 fold increase in proteoglycan content was seen over the low confluent constructs. The improved collagen ratio resulted from a significantly increased type II collagen content and a decreased type I collagen content in over confluent constructs (Fig 2I). Type X collagen (marker of hypertrophic cartilage) gene expression analysis using RT-qPCR demonstrated a significant 3-fold reduction in expression in constructs generated from over confluent BM-MSCs (Fig 2F).

It is clear therefore that cartilage tissue engineering utilising over confluent BM-BMCs produced significantly superior cartilage constructs than those cells harvested at low confluence.

### THE CHONDROGENIC MARKER ROR2 IS DEPENDANT ON CELL DENSITY

The Wnt5a receptor ROR2 has been shown to be predictive of enhanced chondrogenesis in BM-MSCs. Flow cytometry analysis of ROR2 expression in BM-MSCs revealed a direct relationship between culture confluence and ROR2 expression (6). To prove this association 6 individual cell lines were plated at a low cell density (2000 cells/cm^2^) and harvested at serial time points until 24 days in culture for ROR2 analysis using flow cytometry. The gating strategy for ROR2 analysis of these samples is detailed in Supplementary Fig 4A. There is a clear relationship between cell density - as measured by time in culture - and whole population (WP) ROR2 expression (Fig 3A). Cells cultured for 24 days were re-plated (Passage 3) at the initial cell density to assess the effect of re-plating on WP ROR2 expression. Re-plating resulted in an initial decrease in WP ROR2 expression, which was followed by an increase in expression with further days in culture (Fig 3B). This pattern was repeated following further re-plating of cells (Passage 4) cultured for 24 days (Fig 3C). The mean fluorescence index (MFI) indicates the cellular expression of each individual cell interrogated by flow cytometry. The MFI for the fluorochrome APC demonstrated a direct relationship between time in culture and cellular ROR2 expression (Supplementary Fig 4B).

**Figure 4:**
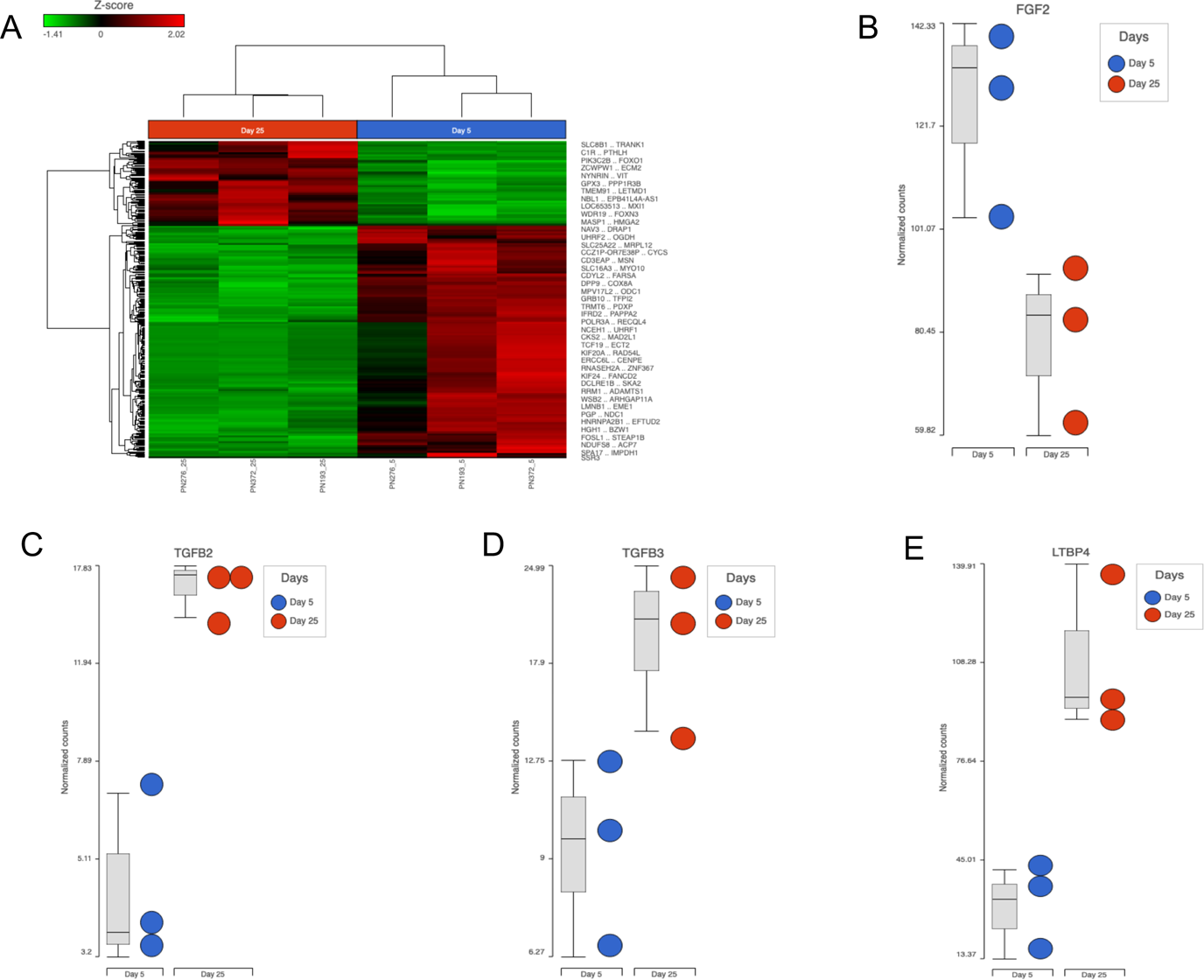
Over Confluent MSC cell lines demonstrate up regulation of the chondrogenic genes - transforming growth factor β2 and β3. A – Hierarchical clustering analysis represented as a heat map. Gene expression was filtered to include those genes with a greater than 2 fold change in expression and an false discovery rate (FDR) of 5%. B – Feature plot demonstrating down-regulation of the growth factor TGF-2 in over confluent cells correlating with the slowing of proliferation once confluence is reached. C-E – Feature plot demonstrating the up regulation (expressed as log fold change) in over confluent BM-MSCs of TGF-beta 2 (3.61 FC; FDR = 0.04), TGF-beta 3 (3.68 FC; FDR = 0.04) and LTBP4 (3.72 FC; FDR = 0.05), key genes involved in chondrogenesis of BM-MSCs. FC = fold change.

Increasing cell density with days in culture is associated with an increase in WP ROR2 expression in BM-MSCs; both have been shown to be markers of enhanced cartilage tissue engineering. It is worth comparing the relationship between these markers and the biochemical composition of the tissue engineered cartilage constructs produced. Cell density - measured at the time of tissue engineering – demonstrated strong positive correlation with improved TII/TI collagen ratio (Fig 3D). On the other hand, the WP ROR2 expression – measured at the time of tissue engineering – showed a weak positive correlation with the TII/TI collagen ratio. The variability demonstrated in the WP ROR2 analysis compared to cell density suggests that cell density may be a more reliable marker of increased TII/TI ratio in tissue engineered cartilage constructs (Fig3D). Correlation of WP ROR2 and cell density with proteoglycan content of the constructs revealed both to have a weakly positive correlation with large variability in data distribution (Fig3E).

### OVER CONFLUENT MSC CELL LINES DISPLAY UP REGULATION OF CRITICAL CHONDROGENIC MARKERS

RNA sequencing provides insights into the gene expression and ultimately the biological activities of cells (15). RNA-Seq analysis was performed on 3 individual BM-MSC cell lines under low and over confluent conditions to assess the gene expression profile of these cells. Preliminary analysis revealed good homogeneity between samples and excellent quality of RNA-seq data for analysis (see Supplementary Figure 5A). Hierarchical analysis was performed after filtering for significance (2-fold change and FDR of 0.05) and displayed as a heat map (see Figure 4A). This demonstrates that genes were predominantly down-regulated in the over confluent cell lines when compared to the same cell lines at low confluence.

Functional analysis revealed down regulation of genes associated with the cell cycle, cellular senescence and regulation of actin cytoskeleton (see Figure 4B). Down-regulation of cell cycle genes points to a cell line slowing proliferation due to the attainment of confluence and limited space to proliferate. This is further corroborated by a significant downregulation of the proliferative growth factor FGF-2 (see Fig 4B). These cells appear to be resilient to cellular stressors such as oxidative stress and DNA damage as demonstrated by the down-regulation of the cellular senescence pathways (16). A slow of proliferation following the formation of condensations is said to occur as these cells adapt to a high cell density environment before differentiation begins (17). Slowing of proliferation as confluence is achieved is reflected *in vitro* in these cells as set out above (see Figure 1B). Murine embryonic stem cells have been shown to differentiate into chondrocytes by the addition of an actin cytoskeleton inhibitor (18).

Furthermore, overexpression of actin binding protein results in upregulation of hypertrophic markers (19). Down-regulation of actin cytoskeleton in over confluent cells may be an indicator of maintenance of a non-hypetrophic phenotype in these MSCs which is in line with our qPCR data (Fig 2F).

Of the genes that were up-regulated in the over confluent cell lines, a few provided possible mechanistic explanations for the improved chondrogenic potential seen in over confluent cell lines. Members of the TGF group of growth factors are known to be essential to the chondrogenic potential of MSCs both *in vitro* and *in vivo* (20–22). TGF-β2 and TGF-β3 are significantly up-regulated in over confluent cell lines (see Figure 4C). Evidence in favour of over confluence cells mimicking condensation formation is provided when considering that TGFβ2 and 3 are upregulated by MCSs immediately after high density pellet generation (23): an *in vitro* design to mimic cellular condensations that occur *in vivo.* Latent TGFβ-binding proteins (LTBP) are components of the ECM and covalently bind TGFβs providing a store for these growth factors in the ECM (24). It follows then that the up-regulated TGFβs in over confluent cell cultures are released and stored in the ECM bound to LTBP in anticipation of chondrogenic differentiation.

Culturing BM-MSCs to over confluence seems to generate an environment akin to cellular condensations in the limb bud prior to chondrogenesis; harvesting these cells at this stage for cartilage tissue engineering provides optimal conditions to improve chondrogenesis.

## MATERIALS & METHODS

### COLLECTION OF BONE MARROW

Adult human BM samples were harvested from patients undergoing elective and emergency orthopaedic surgery at a single institution. Patients provided written informed consent for the harvest of BM and the use of the cells for research in accordance with both the Southmead Hospital Research Ethics Committee (Ref 078/01) and the National Research Authority Committee of the South West (REF 14/SW/1083).

#### Isolation, culture, cryopreservation & thawing of bone marrow MSC’s

Cell culture procedures were carried out using aseptic techniques in a sterile class II laminar flow cabinet (Thermo Scientific, Waltham, USA). Adult BM samples were plated directly into T175 cm^2^ culture flasks (Corning, Appleton Woods, Birmingham, UK) containing adult expansion medium supplemented with 5 ng/mL rhFGF-2 to enhance MSC proliferation.

Approximately 1 mL of bone marrow aspirate was added to each culture flask. The flasks were incubated at 37°C in a humidified incubator containing 5% CO_2_. The MSCs were isolated by adherence to plastic and non-adherent cells were washed away at each media change. The initial media change occurred at 5 days but subsequent media was changed 3 times a week until the cells were approximately 80% confluent. The cells were harvested and frozen in freezing media and stored in a liquid nitrogen facility until further use.

The confluent cells were harvested by aspirating the expansion medium from the culture dish and washed with phosphate buffered saline (PBS). The cells were detached from the culture plastic by adding an appropriate volume of 0.25% trypsin (2.5 g/l) and ethylenediaminetetraacetic acid (EDTA) (0.2 g/l) onto the cells and placed in a 37°C/5% CO_2_ incubator until the cells have detached (≈5 minutes). Once detached the trypsin/EDTA was inhibited by adding double the volume of expansion medium to the flask. The cell suspension was transferred into a 15 mL or 50mL polypropylene tube (Corning) and centrifuged at 1500 revolutions (rpm) for 5 minutes at room temperature. The supernatant was removed and the cell pellet was re-suspended in a known volume and the cells counted using a haemocytometer. The appropriate amount of cells was transferred to a polypropylene tube and centrifuged. The supernatant was removed and the cell pellet was gently re-suspended in freezing medium (comprising two-thirds 10% DMSO in foetal bovine serum (FBS) and one-third MSC expansion medium) at a concentration of 1 x 10^6^ adult MSCs per millilitre. The cryovials (Nalgene, Sigma, Dorset, UK) containing the cells were contained in a Mr Frosty^®^ freezing container (Nalgene) to allow incremental cooling at a rate of 1°C per minute in a −80°C freezer for 12-24 hours before being transferred to a liquid nitrogen storage facility.

Previously frozen cells were collected from liquid nitrogen storage and allowed to thaw in a 37°C water bath. Once thawed 50 000 MSCs (50μL) were seeded into T25 cm^2^ cell culture flasks with the remaining cells (950µL) seeded into T175 cm^2^ cell culture flasks (both Corning) containing the appropriate amount of expansion medium supplemented with 5 ng/mL rhFGF-2. The cells were maintained in a humidified incubator containing 5% CO_2_ at 37°C. Media was renewed three times a week until approximately 80% confluence was reached.

The cells were cultured in monolayer in T25, T75 and T175 cm^2^ cell culture flasks (all from Corning) depending on the cell number or culture conditions for each experiment. The cells were passaged once they reached 80-90% confluence. The cells were harvested, counted and centrifuged as outlined above. Once counted the appropriate concentration of cells was re-plated in culture flasks in expansion medium supplemented with 5 ng/mL rhFGF-2 and maintained in a humidified incubator containing 5% CO_2_.

### BM-MSC CHARACTERISATION

#### Growth Kinetics

BM-MSC cell lines from 5 individual donors from passage 2 were plated at a cell density of 2000 cells/cm^2^ and permitted to proliferate in expansion media supplemented with rh-FGF. At defined time points the cells were harvested, counted and analysed using flow cytometry to determine the cell density at harvest the population doubling time and the cell viability expressed as a percentage of the WP at flow cytometry.

The population doubling time measures the time taken for the cell population to double in number as is measured by the following equation: PDt = time in culture x (log 2/log (number of cells at passage/ number of cells plated)

Flow cytometry was performed in order to qualtify the celluar viability using propidium iodide as a viability marker. The cells were harvested, counted and centrifuged and washed twice in flow buffer −0.5% mass/volume (w/v) bovine serum albumin (BSA) in PBS (both from Sigma). A total of 50 000 cells from each sample were transferred into a sterile polystyrene test tube (BD Bioscience, Oxford, UK). Propidium iodide (Sigma) was used to identify dead cells, 2.5 µg/mL was added immediately prior to analysis. A minimum of 10 000 single live events was used as a standard cell count in each analysis. Acquisition and analysis was performed on a MACSQuant Analyzer 10 flow cytometer (Metenyi). Final analysis was performed using FloJo Software (Treestar, Ashland, USA).

#### Adipogenic and Osteogenic potential of OA and Trauma BM-MSCs

Six individual cell lines cultured to low and over confluence were stimulated to differentiate along adipogenic and osteogenic lines in monolayer culture. The resultant culture was analysed histologically with the appropriate stains.

The cultured BM-MSCs at passage 3 were harvested, counted and centrifuged once 80% confluency was reached. A total of 3.7×10^4^ cells were added to the middle wells of a 24-well plate (Appleton Woods) for each cell line. The remaining wells were filled with sterile water containing antifungal agents. Once the cells were adherent the cells undergoing differentiation were cultured in basal medium supplemented with 1% v/v Adipogenic Supplement (R&D Systems) while the control cells not undergoing differentiation were cultured in basal medium. Basal medium contains: Dulbecco’s modified Eagle’s medium (DMEM) supplemented with 4500 mg/L glucose, 100 units/mL penicillin, 100 µg/mL streptomycin, 1nM sodium pyruvate, 80 µM ascorbic acid-2-phosphate, 100 nM dexamethasone (all from Sigma), 2 mM glutamax and 1% v/v 100x insulin transferrin selenium (ITS) (both from Invitrogen). The cells were incubated in a humidified incubator with 5% CO_2_ at 37°C and media renewed every 3 days for 21 days.

Following 21 days of adipogenic stimulation, the cells were washed in sterile PBS and fixed in 4% v/v paraformaldehyde for 30 minutes at room temperature. The fixative was removed and the cells were washed in 60% isopropanol. To visualise any lipid vacuoles, 0.3 % Oil red O (Sigma) was used to stain the cells and was left for 30 minutes at room temperature. The stained cells were photographed using bright field microscopy.

The same cultured BM-MSCs as above were harvested, counted and centrifuged. A total of 7.4×10^3^ cells from each cell line were added to the middle wells of a 24-well plate (Appleton Woods) with the remaining wells containing sterile antifungal water. Once adherent, the cells undergoing differentiation were cultured in basal medium supplemented with 5% v/v Osteogenic Supplement (R&D Systems) with the control cells cultured in basal medium. The samples were incubated in a humidified incubator with 5% CO_2_ at 37°C with media renewed every 3 days for 14 days.

Following 21 days of osteogenic differentiation, the cells were washed in PBS and fixed with 70% v/v ice-cold ethanol for 1 hour at 4°C. The fixative was carefully removed and the cells washed in PBS. The ECM was stained with 40 mM alizarin red (Sigma) for 5 minutes at room temperature. The stain was removed and the cells washed with PBS to remove the stain.

### CHONDROGENIC POTENTIAL OF OA AND TRAUMA BM-MSCS

BM-MSCs from 10 individual cell lines were permitted to proliferate in expansion medium for 5 days (low confluence) and 24 days (over confluence)before bwing harvested and stimulated to undergo chondrogenic differentiation. The 3D cartilage tissue engineering constructs were used for gene, biochemical and histological analysis.

#### Three-Dimensional Cartilage Tissue Engineering

Ethylene oxide was used to sterilise the PGA scaffolds (Anderson Caledonia, Bellshill, Scotland). In order to remove contaminants, the scaffolds were immersed in absolute ethanol for 5 minutes and washed three times in PBS. The scaffolds were coated with 10 µg/mL of fibronectin (Sigma) and allowed to dry overnight in a sterile laminar flow cabinet. To prevent the cells from adhering to the culture plate, the required wells were coated with 500 µL 1% agarose. The dried fibronectin-coated scaffolds were added to the agarose wells and rehydrated with expansion medium. The MSCs were trypsinised, counted and centrifuged. A total of 5×10^5^ cells per scaffold were re-suspended in 30 µL of expansion medium. The expansion medium was removed from the scaffolds immediately prior to seeding. The cells were carefully seeded onto the scaffolds taking care to equally distribute the cell suspension. The scaffolds were placed in a humidified incubator at 37°C containing 5% CO_2_ overnight to allow the cells to adhere to the scaffold. The scaffolds were then turned and incubated for a further 2 hours to allow even cell distribution and adhesion; the scaffolds were then supplied with ITS basal medium supplemented with 10 ng/mL rhTGFß3 (R&D Systems), 100 nM dexamethasone, 80 µM ascorbic acid 2-phosphate (both Sigma) and incubated on a rotating platform at 50 rpm in a humidified incubator with 5% CO_2_ at 37°C. Media was renewed every 3 days for a total of 35 days; insulin at 10 µg/mL concentration was added to the basal medium supplement from day 7. After a total of 35 days in culture the scaffolds were harvested, photographed and placed in pre-weighed tubes. The constructs were stored at −20°C until freeze-drying.

The tissue engineered cartilage constructs were used to analyse the matrix composition in relation to the glycosaminoglycan (GAG), collagen I and collagen II content of these constructs.

Prior to digestion the constructs were lyophilised in a freeze dryer (Savant Modulyo, Thermo Scientific) for 24 hours. The freeze-dried sample were then weighed and subtracted from the tube weight to determine the dry weight of the constructs. The dried constructs were digested by adding trypsin in two steps.

The initial trypsin aliquot was added together with three proteases to prevent non-specific protease activity in the constructs and incubated at 37°C on a rotating platform (100 rpm) for 12 hours. The second aliquot of trypsin was added to the mixture and incubated at 65°C for a further 3 hours; the mixture was subsequently boiled for 15 minutes to inhibit the trypsin activity. The volume of trypsin used was determined by the dry weight of the constructs – 50 µL per 0.5 mg. The resultant mixture was then centrifuged at 13 000 rpm for 5 minutes and the supernatant aspirated and stored at 4°C for analysis. The remaining undigested material was further freeze dried and weighed to determine the final dry weight of the digested ECM: Dry weight of digested ECM = Dry weight of freeze-dried scaffold/ECM construct – Dry weight of undigested material.

#### Gene analysis of tissue engineered cartilage

The cartilage tissue constructs for gene analysis were stored until processing at −80°C. The construct was processed using the Total RNA Purification Kit (Norgen Biotek). The constructs were prepared for processing by cutting each sample into 2mm x 2mm pieces and immediately immersed in the lysis buffer. The sample was crushed using a small pestle and repeatedly passed through a 21 gauge needle until the sample passed through easily. The sample was then passed through a 25 gauge needle to shear the genomic DNA. Absolute ethanol was then added to the lysate and vortexed to ensure adequate mixing.

The lysate was then transferred into a spin-column and centrifuged at room temperature at 14 000 rpm for 2 minutes. The remaining RNA in the spin-column – after discarding the flow-through - was washed three times and centrifuged to leave the RNA collected in the resin thoroughly dry. The bound RNA was then eluted and quantified to assess concentration and purity using a NanoDrop spectrophotometer (Thermo Scientific).

Purified RNA was transcribed into cDNA using the PrimeScript Reagent Kit (Takara). The volume equivalent of 0.2 µg of RNA was reverse transcribed in a 40 µL reaction volume. The total reaction volume consisted of 6 µL 1 x concentration buffer, 6 µL enzyme mix, 25 pmol oligo dT primers, 50 pmol random hexamers and RNase free dH2O. The reaction mixture was allowed to incubate in a thermal cycler at 37°C for 15 minutes to facilitate reverse transcription and at 85°C for 5 seconds to inactivate the enzymes. The cDNA was stored at −20°C until further use.

The expression of the hypertrophic chondrogenic marker gene – collagen X – was measured using RT-qPCR with TaqMan^®^ as the fluorescence reporter. A one-step RT-qPCR reaction will be used to amplify the cDNA using the QuantStudio Real-Time PCR System (ThermoFisher Scientific, Waltham, USA). Each sample will be processed in triplicate with beta actin used as a control gene. Comparitive quantification of the selected chondrogenic genes was calculated using the ΔΔCT_t_ method.

#### Biochemical analysis of cartilage constructs

The proteoglycan content of each scaffold was measured using a colourmetric assay to detect sulphated GAG (25). A set of 11 standards was used for this assay and prepared by diluting chondroitin sulphate in dH_2_O in 5 µg/mL increments from 0 µg/mL to 50 µg/mL. A 96-well, round-bottomed microplate (Corning) was used and set up as follows: a row of blank (B) negative control wells containing 20 µL of dH2O, the 11 standards (S1-11) were added in duplicate with 20 µL being added to each well and finally 20 µL of each sample (U) was transferred to the plate in duplicate. A total of 250 µL of BMB was added to each well and the optical density of the samples was read using a Glomax plate reader set at 525 nm (Promega, Madison, USA). The amount of GAG in each sample is expressed as a percentage of the dry weight of the digested ECM.

The collagen content of tissue engineered cartilage was measured by an inhibition ELISA assay using a primary antibody against a peptide (AH23) specific to collagen type I (26). A primary antibody (AH23-1319) was used to bind type I collagen present in the digested ECM; the remaining antibody not bound to the sample collagen was measured by allowing the remaining antibody to bind to AH23 pre-coated on a separate plate. A secondary antibody conjugated to alkaline phosphatase was added to determine the amount of antibody not bound to the initial sample. The collagen content of the sample ECM can then be calculated and expressed as a percentage of the dry-weight of the digested ECM.

A concentration of 40 µg/mL of the procollagen AH23 peptide was prepared by dissolving the peptide in carbonate buffer. A total of 50 µL of the procollagen solution was then added to the inner wells of an Immulon-2 high binding, flat-bottomed 384-well plate (Thermo Fisher Scientific, Loughborough, UK). The plates were stored at 4°C for 72 hours to allow the procollagen to bind and coat the plates. Prior to use the plates were washed thoroughly with PBS/Tween and dried. The plates were then ‘blocked’ against non-specific binding by coating the wells with 80 µL of 1% BSA/PBS which was left at room temperature for 30 minutes. The plates were thoroughly washed again in PBS/Tween and dried at 50°C for 15 minutes before being stored at 4°C until required.

Uncoated round bottomed 384-well microplates (Corning) were ‘blocked’ with 1% BSA/PBS at room temperature for 30 minutes. These plates were then washed with PBS/Tween and dried at 37°C for 30 minutes. The anti-peptide primary antibody was prepared by diluting 1:1000 in Tris buffer containing 4% (v/v) Triton-X100. The plate was then set up as follows: 18 rows of blank (B) wells containing 170 µL of 0.8% SDS/Tris, 12 rows of maximum binding (MB) wells each containing 85 µL of 0.8% SDS/Tris, nine standards (S1-S9) prepared by double diluting the AH23 peptide in a range from 3.9 ng/mL to 1000 ng/mL in triplicate and finally samples (U) in triplicate after appropriate dilutions in SDS/Tris. The primary antibody was added to all the wells except for the blanks. The plates were tightly sealed and incubated overnight at 37°C. The samples were then carefully transferred into the appropriate well of the pre-coated procollagen plate and incubated at room temperature for 30 minutes. The plates were washed with PBS/Tween and dried thoroughly. The anti-rabbit goat secondary antibody conjugated to alkaline phosphatase was prepared by diluting 1:1000 in 1% BSA/PBS containing Tween-20. This secondary antibody was added to all the wells and incubated at 37°C for 2 hours. The plate was washed thoroughly with PBS/Tween and lastly with dH_2_O. The alkaline phosphate substrate diluted in diethanolamine buffer was added to the wells and allowed to develop for 20-30 minutes. The absorbance was read at 405 nm using a Mithras plate reader (Berthold Technologies, Zug, Switzerland). The quantity of collagen type I was expressed as a percentage of the dry weight of digested ECM.

The collagen type II content of the tissue engineered cartilage will be measured using an inhibition ELISA by using a primary antibody (COL2-3/4) against a peptide specific to collagen type II (CB11B) (27).

The collagen peptide CB11B of concentration 60 µL/mL was prepared by diluting in carbonate buffer. The collagen peptide was denatured by heating for 20 minutes at 80°C. A total of 40 µL of the denatured collagen peptide was added to the inner wells of a high binding, flat-bottomed 384-well plate (Corning). The plates were then wrapped in cling-film and incubated at 4°C for 72 hours to ensure adequate coating of the collagen to the plates. Following this, the plates were washed thoroughly with PBS/Tween and dried. To prevent non-specific binding the plates were ‘blocked’ by placing 50 µL of a 1% BSA/PBS solution into each well and incubated at room temperature for 30 minutes. The plates were again washed with PBS/Tween and dried for 15 minutes at 50°C and stored at 4°C until required.

Prior to use, low binding, flat bottomed 384-well microplates (Corning) were ‘blocked’ to prevent non-specific binding with 50 µL 1% BSA/PBS and incubated at room temperature for 30 minutes. The plates were then washed thoroughly in PBS/Tween and dried. Seven standards were made from CB11B in concentrations ranging from 0.5 µg/mL to 6 µg/mL made up by double diluting in Tris buffer and the COL2-3/4m primary antibody was prepared by diluting at a ratio of 1:600 in Tris buffer. The plates were set up as follows: 24 blank (B) negative control wells containing 40 µL 50 mM Tris buffer, 18 maximum binding (MB) positive control wells containing 20 µL 50 mM Tris buffer, 20 µL of the 7 standards (S1-7) in triplicate and finally samples (U) in triplicate diluted to the appropriate concentration in Tris buffer. In all the wells except the blanks, 20 µL of antibody was added. The plates were sealed and incubated at 37°C overnight. Following incubation, 10 µL was transferred into the corresponding well of the previously collagen-coated 384-well plate. The plate was sealed and incubated at room temperature for exactly 30 minutes. The plates were then washed three times in PBS/Tween and dried at 37°C for 30 minutes. The goat anti-mouse secondary antibody was prepared by diluting 1:1000 in 1% BSA/PBS containing Tween-20 and 10 µL was added to each well following drying. The plate was sealed again and incubated at 37°C for 2 hours. After the incubation period, the plates were washed three times in PBS/Tween and once in dH_2_O. Alkaline phosphate substrate in diethanolamine buffer was added to each well of the plate and allowed to develop at 37°C for 10 minutes. The plate’s absorbance was then read at 405 nm on a Mithras plate reader (Berthold). The quantity of the collagen type II was determined from the standard curve and expressed as a percentage of the dry weight of digested ECM.

#### Histological analysis of tissue engineered cartilage

The tissue engineered cartilage constructs were immersed in OCT embedding matrix (BDH Chemicals, Poole, UK) and suspended over liquid nitrogen until the OCT solidified thus embedding the constructs in OCT. These were then cut into 7 μm sections using a cryostat before being transferred to polysine microscope slides. The sections were permitted to dry at room temperature for 30 minutes and then fixed in 4% (w/v) paraformaldehyde (Sigma) at 4°C for 30 minutes. The sections were then washed in PBS and stored until staining.

Prepared sections were rehydrated in dH_2_O for approximately 5 minutes. Haematoxylin (Vector Laboratories, Peterborough, UK) was initially used to stain the sections. After approximately 2 minutes any access haematoxylin stain was removed with dH_2_O before 1% eosin (Vector Laboratories) was added. After 2 minutes the sections were rinsed in 70% then 90% and then 100% ethanol before being cleared in xylene for 5 minutes. The sections were then mounted in DPX mounting medium and covered with a cover slip.

The sections were rehydrated as above and stained with 0.02% (w/v) fast green (Sigma Aldrich) for 4 minutes and then rinsed in 1% acetic acid. The 0.1% (w/v) Safranin-O stain (Sigma Aldrich) was then added to the sections for 6 minutes. The sections were then rinsed in 70%, 90% and 100% ethanol before being cleared in xylene for 5 minutes. The sections were then mounted in DPX mounting medium and covered with a cover slip.

The prepared sections were subjected to enzymatic retrieval by initially treating with a 0.1% (w/v) pronase (Sigma Aldrich) solution before being incubated at room temperature in a humidified chamber. The sections were then washed with PBS and then treated with 2% (w/v) hyaluronidase (Sigma Aldrich) at 37°C for 30 minutes in a humidified chamber. Following this the sections were washed twice in PBS for 2 minutes each and treated with 3% hydrogen peroxidase at room temperature for 10 minutes and washed in PBS.

Prior to staining for collagen type II the sections were blocked with Novolink protein (Leica Biosystems, Weltzlar, Germany) for 5 minutes at room temperature. After a rinse in PBS the primary collagen type II antibody - diluted to 1 in 4000 in Tris buffered saline (TBS) solution (20 mM Tris and 150 mM NACl) and 1% goat serum – was incubated overnight at 4°C with an appropriate isotype control. The primary antibody to collagen type II used was II-II6B3 (DSHB, deposited by Linsenmayer, T.F.) and the isotype control was a mouse IgG1 istotype (2b Scientific, UK). The antibodies were washed in TBS three times for three minutes at a time before the Novolink post primary block (Leica biosystems) was applied for 30 minutes at room temperature. After another three washes in TBS the Novolink polymer (Leica biosystmes) was applied for another 30 minutes at room temperature. The sections were then treated with DAB (3,3’-Diaminobenzidine) and incubated at room temperature for 3 minutes. The DAB was removed with dH2O and haematoxylin stain was applied for 3 minutes before a final wash in preparation for dehydration. The sections were dehydrated in 70%, 90% and 100% ethanol for 5 minutes each and cleared in xylene and mounted using DPX and covered with a cover slip.

The sections were initially subjected to enzymatic retrieval. Prior to staining the sections were blocked with Novolink protein (Leica Biosystems) for 5 minutes at room temperature. The primary collagen type I antibody (ab23446, Abcam, Cambridge, UK) – diluted 1 in 100 in TBS with 1 % goat serum – was incubated overnight at 4°C with the appropriate isotype control (ab170190, Abcam). Following incubation, the sections were washed three times in TBS. The final preparation of the slides is as was performed in the preparation of the collagen type II stained slides as above.

### ROR2 IMMUNOPHENOTYPING BY FLOW CYTOMETRY

#### Analysis of ROR2 in cultured adult BM-MSCs

Bone marrow derived MSCs were harvested, counted, centrifuged and washed twice in flow buffer (0.5% w/v BSA in PBS (both from Sigma)). For each sample 200 000 cells was transferred into sterile polystyrene test tubes (BD Bioscience) and stained with the appropriate primary antibody (see Table 1). The samples were then incubated at −4°C in the dark for 30 minutes and then washed twice with flow buffer. The secondary antibody was introduced and again allowed to incubate at −4°C in the dark for 30 minutes. After removing the unbound secondary antibodies with flow buffer, propidium iodide (Sigma) was used to identify dead cells: 2.5 µg/mL was added immediately prior to analysis. The cells were processed using the MACSQuant Analyser 10 flow cytometer (Mitenyi) and the data analysed with FloJo software (Treestar).

**Table 1:**
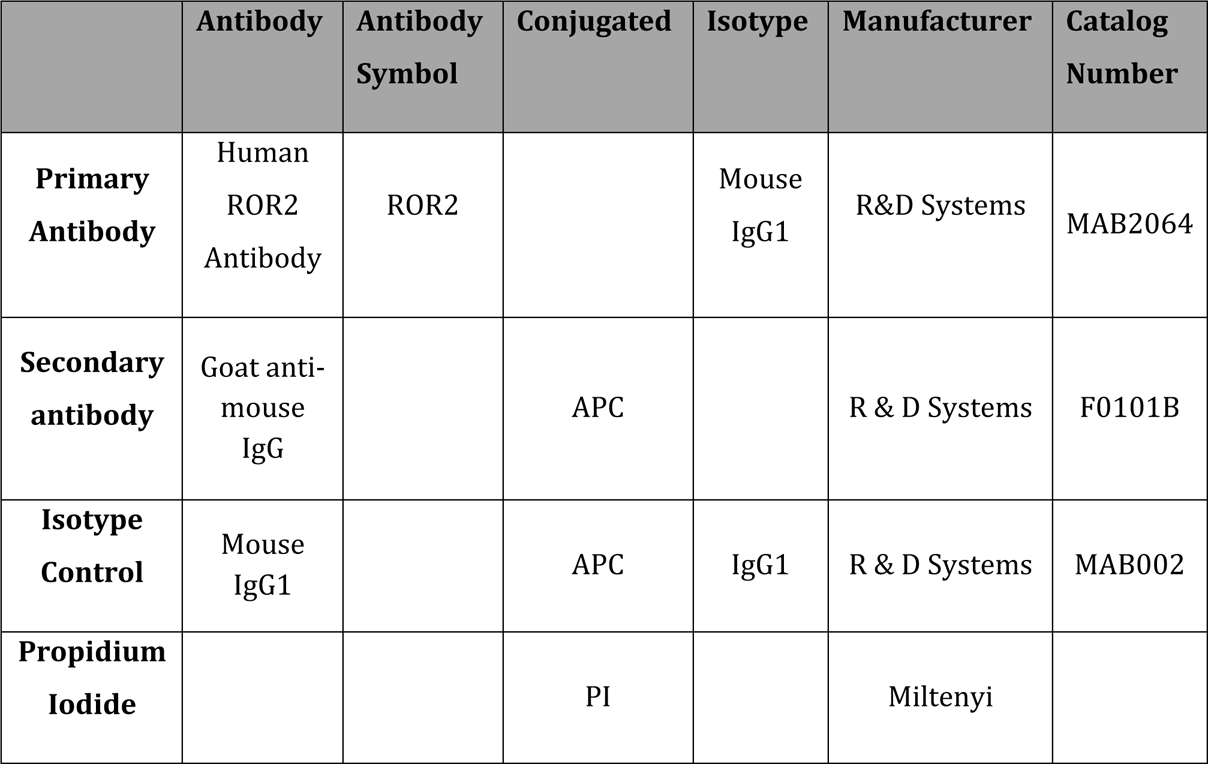
Antibody panel used in flow cytometry analysis of ROR2 expression in BM-MSCs from OA and trauma patients

|  | Antibody | Antibody Symbol | Conjugated | Isotype | Manufacturer | Catalog Number |
| --- | --- | --- | --- | --- | --- | --- |
| **Primary Antibody** | Human ROR2 Antibody | ROR2 |  | Mouse IgG1 | R&D Systems | MAB2064 |
| **Secondary antibody** | Goat anti-mouse IgG |  | APC |  | R & D Systems | F0101B |
| **Isotype Control** | Mouse IgG1 |  | APC | IgG1 | R & D Systems | MAB002 |
| **Propidium Iodide** |  |  | PI |  | Miltenyi |  |

**Table 2:** Genetic Pathway Analysis comparing low to over confluent cell lines

| KEGG Pathway | Enrichment Score | p Value |
| --- | --- | --- |
| Cell Cycle | 17.24 | 3.27E-8 |
| Cellular Senescence | 6.48 | 1.54E-3 |
| Regulation of Actin Cytoskeleton | 3.95 | 0.02 |

#### The effect of confluence/cell density on ROR2 expression

Cells from 5 different BM-MSC samples were thawed and plated at a cell density of 2666 cells/cm^2^ in a T75 cm^2^ culture flask (Corning). Once the cells reached 80-90% confluence they were passaged and re-plated at a cell density of 2000 cells/cm^2^ in T25 cm^2^ culture flasks (Corning). For each cell line a total of seven T25 cm^2^ culture flasks were required for flow cytometry analysis. To assess the impact of confluence on ROR2 expression, flow cytometry was performed at day 3, 6, 10, 14, 17, 21 and 24 post plating. The procedure for immunophenotyping these cells is described above. At each interval, prior to analysis, the cells were photographed using bright field microscopy.

#### The effect of re-plating on ROR2 Expression

Utilising the same 5 samples as above, following harvesting of the cells cultured for 24 days these BM-MSCs were again plated at a cell density of 2000 cells/cm^2^ in T25 cm^2^ culture flasks (Corning) and allowed to proliferate. These cells were analysed by flow cytometry on day 1, 2, 3, 7, 14 and 24 to assess the effect of re-plating on ROR2 expression. In order to assess the effect over multiple passages the process was repeated for a further passage.

#### ROR2 expression prior to cartilage tissue engineering

Prior to tissue engineering, low confluent (5 days in culture) and over confluent (24 days in culture) BM-MSCs from 10 paired individual cell lines were analysed using flow cytometry to determine the WP ROR2 positivity prior to tissue engineering. The processes for 3D tissue engineering and flow cytometry analysis has been outlines above.

### RNA-SEQ ANALYSIS OF LOW AND OVER CONFLUENT BM-MSCS

Adult BM-MSCs from 3 individual cell lines were thawed and plated in proliferation media. Once 80% confluency was reached the cells were harvested and plated at a cell density of 2000 cells/cm^3^ and permitted to proliferate for 5 days (low confluence) and 24 days (over confluence) before being harvested for RNA extraction.

The construct was processed using the Ambion Purelink RNA Purification Kit (ThermoFisher Scientific). The sample was then passed through a 21 gauge needle to shear the genomic DNA. Absolute ethanol was then added to the lysate and vortexed to ensure adequate mixing. The lysate was then transferred into a spin-column and centrifuged at room temperature at 14 000 rpm for 2 minutes. The remaining RNA in the spin-column – after discarding the flow-through - was washed three times and centrifuged to leave the RNA collected in the resin thoroughly dry. The bound RNA was then eluted and quantified to assess concentration and purity using a NanoDrop spectrophotometer (Thermo Scientific). The RNA was then stored in lysis buffer (with B-MB) in RNA-free centrifuge tubes and stored at −80°C and shipped to Novogene (Beijing, China) for medical transcriptome analysis. The data was analysed using Partek^®^ Flow^®^ software.

---

**Supplementary Figure 1:**
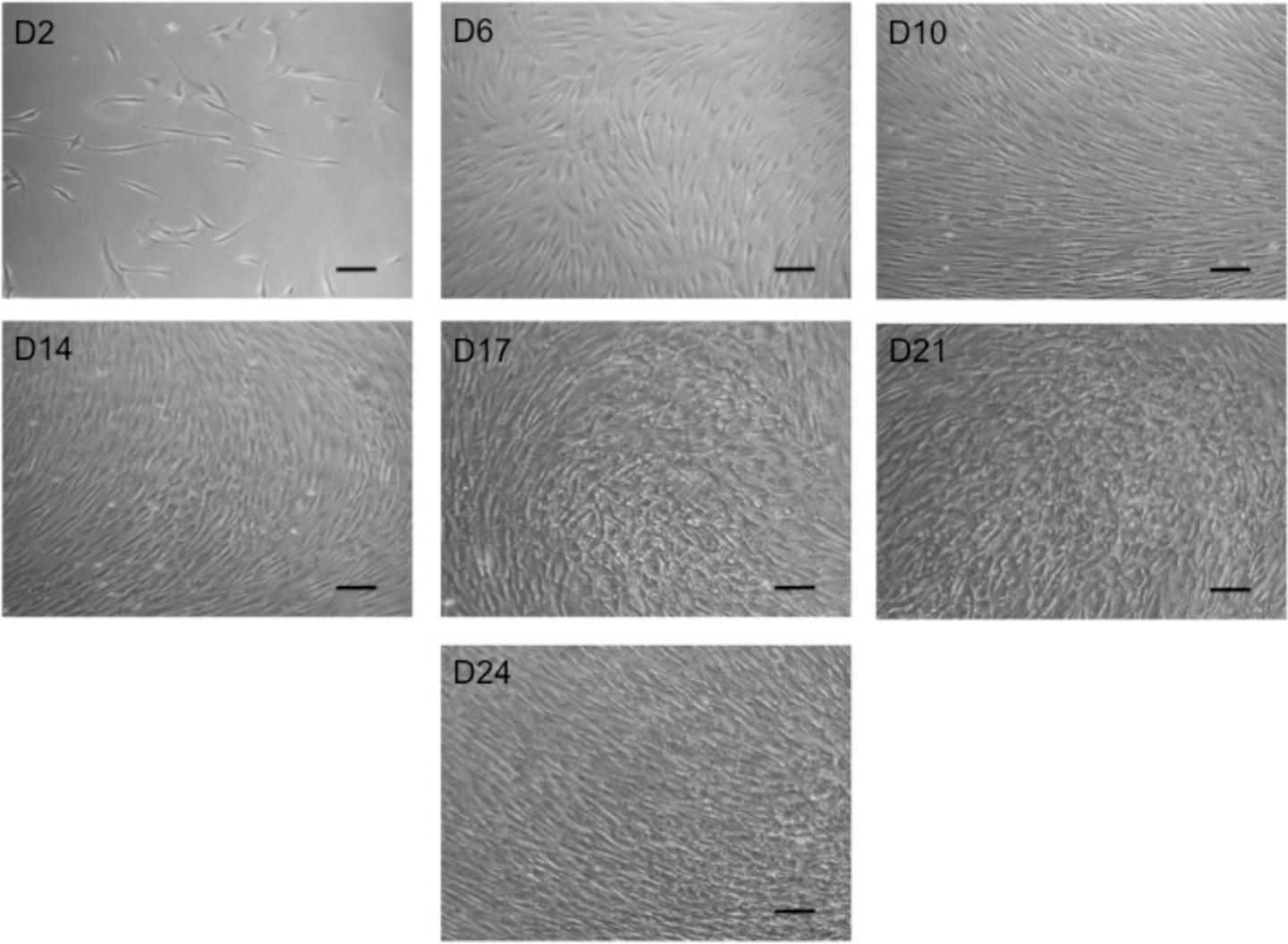
Visual assessment of confluent cells over time. Representative bright field microscopy images of BM-MSCs plated at 2000 cells/cm^2^ and cultured for 24 days in expansion media before being photographed at specified time points. Scale bar corresponds to 100 μm. Original magnification 10x.

**Supplementary Figure 2:**
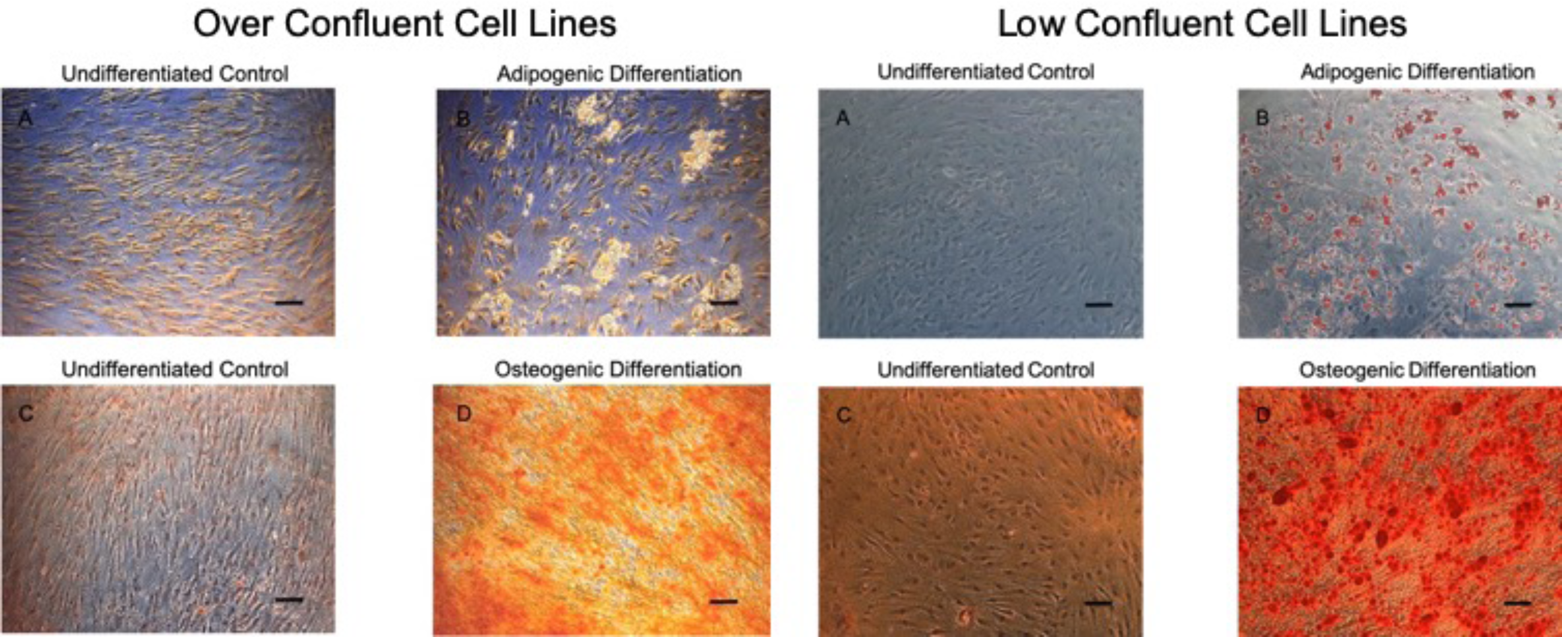
Multipotency of over confluent BM-MSCs. Over Confluent Cell Lines - The potential of BM-MSCs cultured to over confluence was assessed by stimulating 6 individual cell lines (initially cultured for 24 days in proliferation media) with adipogenic (A & B) and osteogenic (C & D) differentiation media. The monolayer cultures were differentiated in triplicate and made use of controls containing no differentiation media. Following 21 days of differentiation the monolayer cultures were stained with the appropriate stains and photographed using bright field microscopy. Representative images of a single cell line stained with Oil Red O to demonstrate lipid vacuoles in an undifferentiated control (A) and differentiated sample (B) as well as those stained with Alizarin red demonstrating calcium deposits in control (C) and differentiated samples (D). Low Confluent Cell Lines – The multipotency of the same 6 cell lines were stimulated to undergo adipogenic (A-B) and osteogenic (C-D) differentiation following 8 days of cell culture. Following 21 days of differentiation the monolayer cultures were stained with the appropriate stains and photographed using bright field microscopy. Representative images of a single cell line stained with Oil Red O to demonstrate lipid vacuoles in an undifferentiated control (A) and differentiated sample (B) as well as those stained with Alizarin red demonstrating calcium deposits in control (C) and differentiated samples (D). Scale bar corresponds to 100 μm. Original magnification 10x.

**Supplementary Figure 1:**
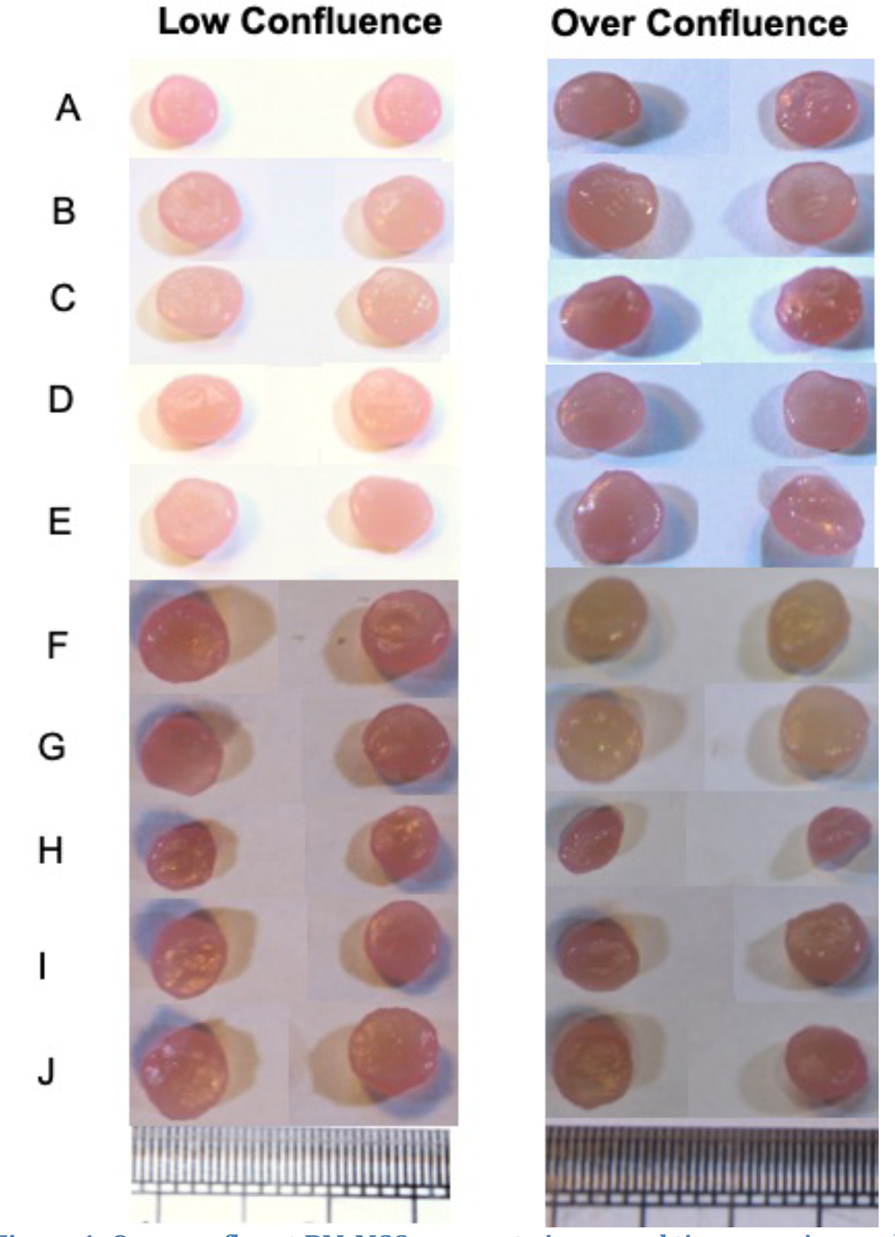
Over confluent BM-MSCs generate improved tissue-engineered cartilage. A – Macroscopic images of paired three-dimensional cartilage tissue engineering using PGA scaffolds coated in fibronectin and generated (in duplicate) using 10 individual BM-MSCs harvested at low and over confluence and cultured for 35 days in the presence of TGF-β3. Scale bar represents 1mm. C - Duplicate three-dimensional tissue engineered cartilage scaffolds generated from paired BM-MSCs from low confluent and over confluent states were freeze dried, digested in trypsin and analysed for the presence of type II collagen and type I collagen. Each point represents the mean quantity of collagen type II for each duplicate sample and the line demonstrates the population mean. Statistical analysis was performed using a paired t test with a p value of <0.05 considered statistically significant. ** = p<0.01.

**Supplementary Figure 2:**
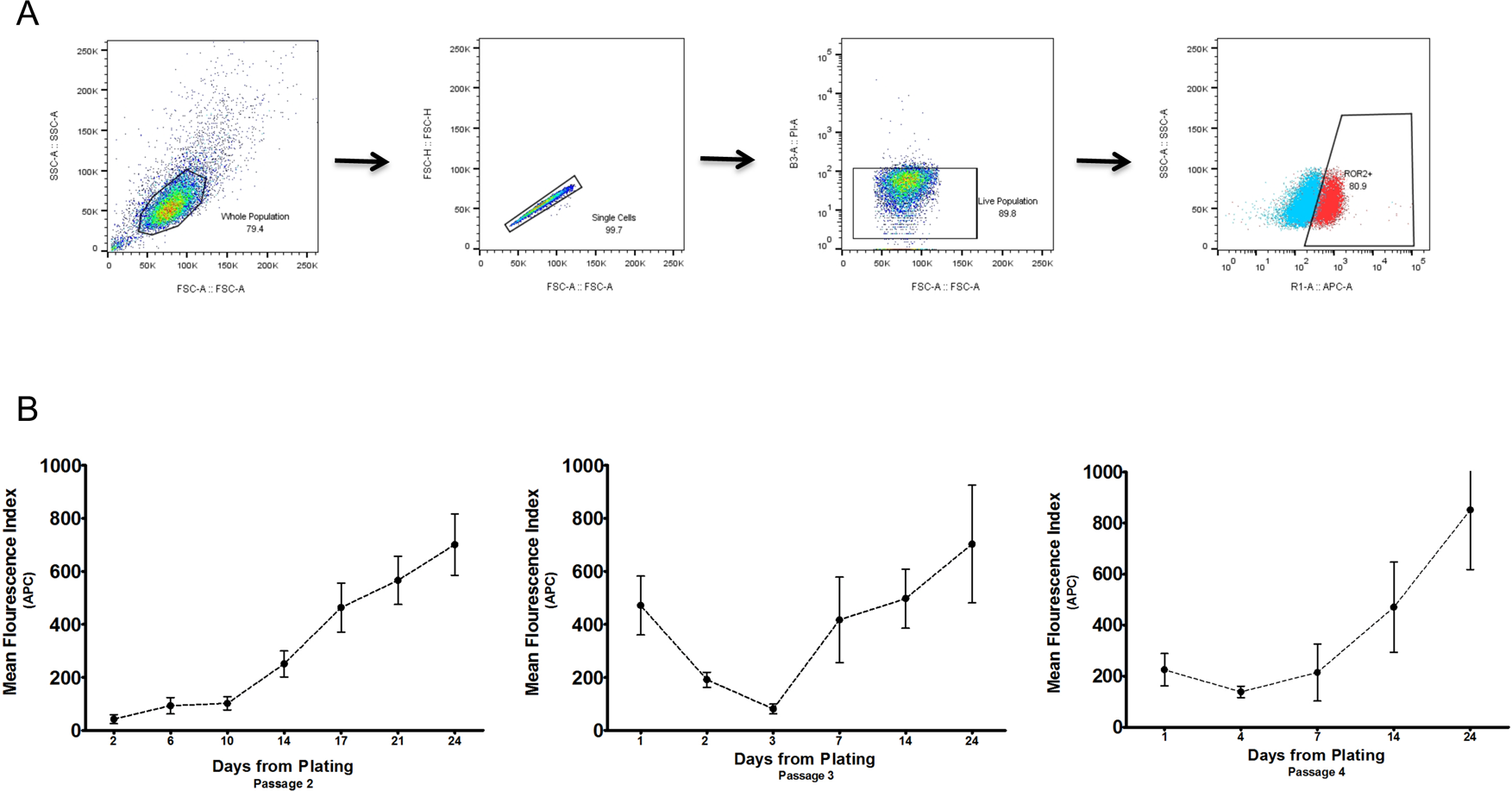
The chondrogenic marker ROR2 is dependant on cell density. A – The flow cytometry gating strategy for WP ROR2 analysis is demonstrated. B -. BM-MSCs from 5 individual cell lines were plated at a cell density of 2000 cells/cm^2^ and permitted to proliferate in expansion media before being harvested, counted and interrogated using flow cytometry to determine the mean fluorescence index for the fluorochrome APC at defined time points until 24 days in culture. Following analysis of cells at 24 days post plating these cells were then re=plated at a cell density of 2000 cells/cm^2^ and permitted to proliferate and again analysed for the mean fluorescence index at defined time points. The process was repeated for a further passage.

**Supplementary Figure 5:**
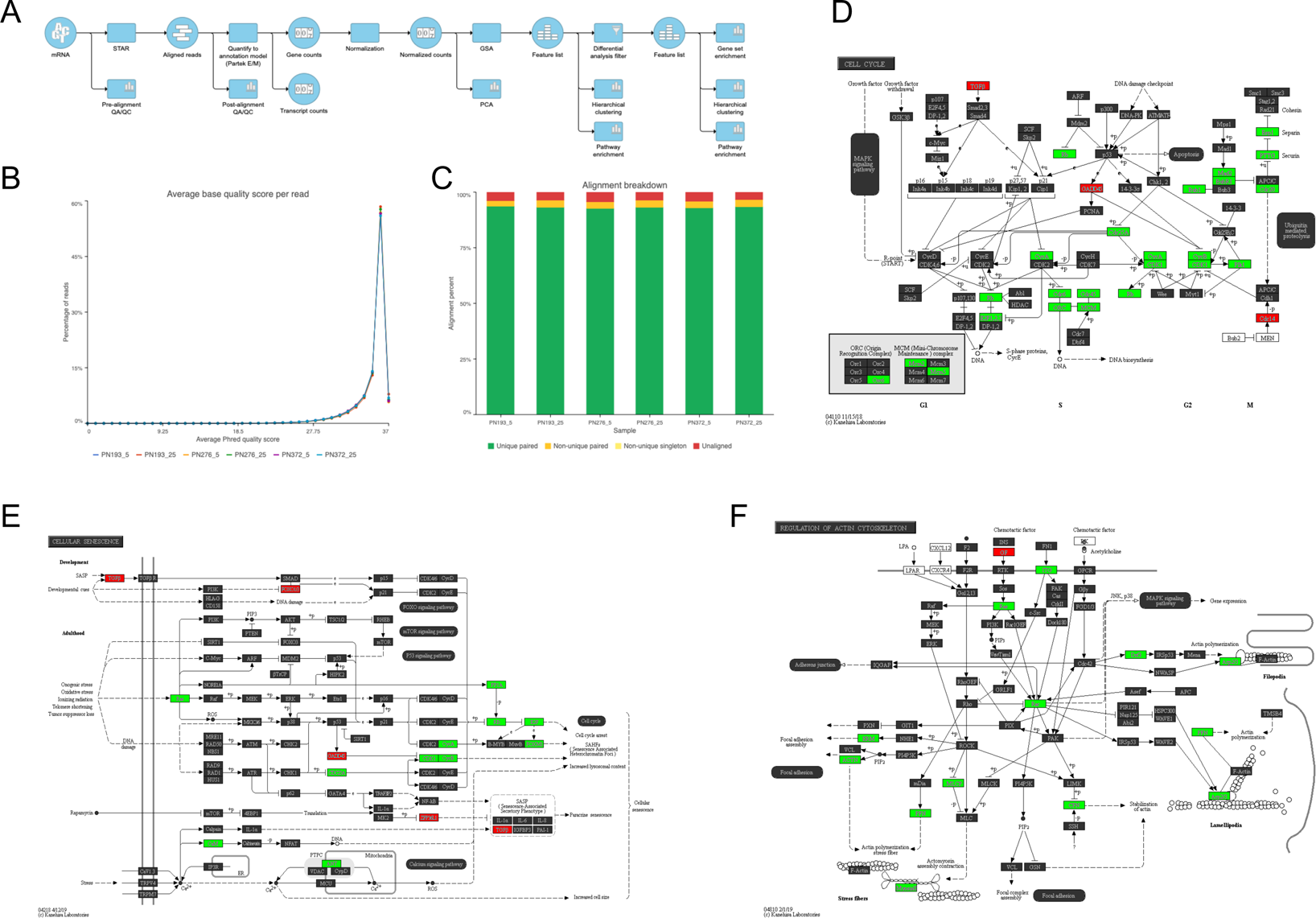
RNA-Seq analysis comparing gene expression of low and over confluent BM-MSC cell lines. A - RNA sequencing was performed on 3 individual cell lines cultured to low confluence (5 days) and over confluence (25 days). Gene specific analysis was performed using Partek^®^ Flow^®^ software utilising the analysis pipeline as shown. B-Pre-alignment QA/QC analysis demonstrated an average read quality in excess of 35 across all 6 samples. C – Following alignment using STAR a post alignment QA/QC analysis was performed which demonstrated alignment in excess of 95% across all samples. D, E&F – Pathway enrichment was performed utilising the KEGG database with pathways for cell cycle (D), cellular senescence (E) and regulation of actin cytoskeleton (F).

**Supplementary Table 1:**
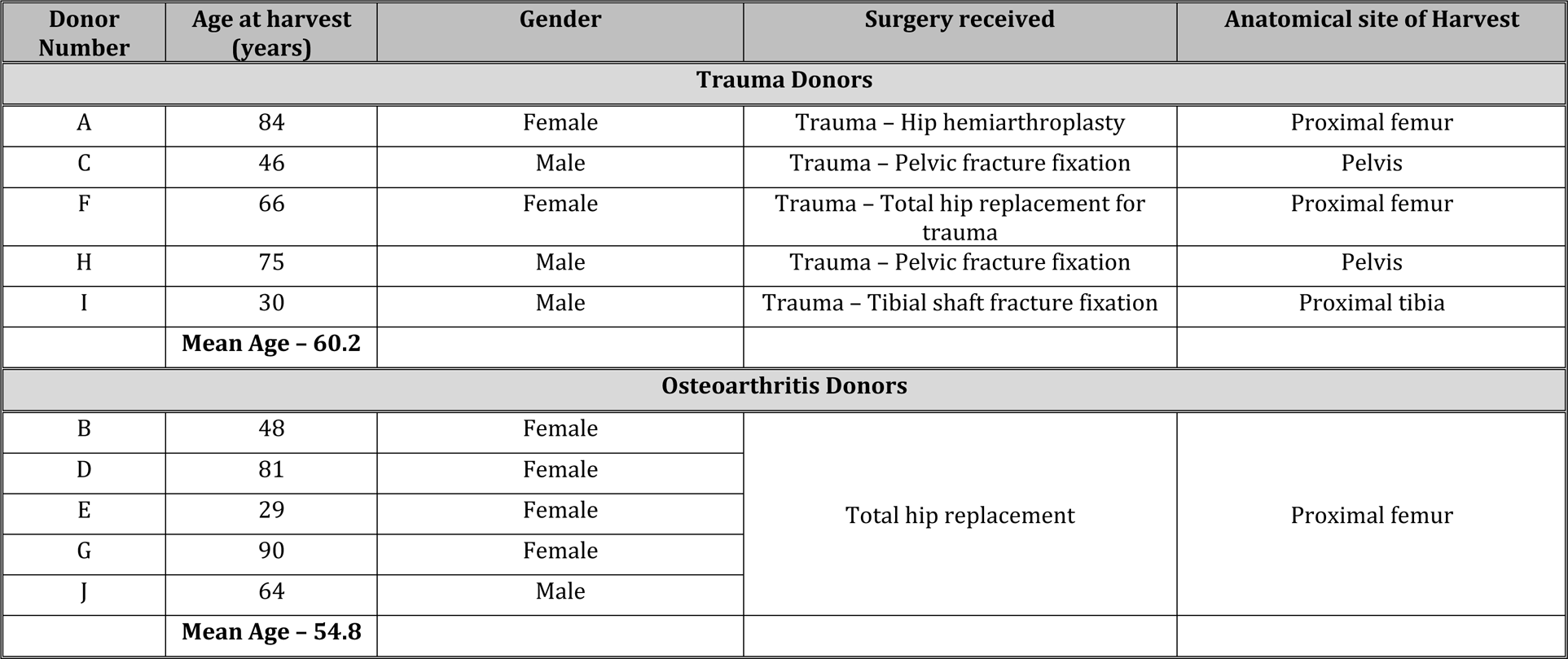
Age and characteristics of BM donors

| Donor Number | Age at harvest (years) | Gender | Surgery received | Anatomical site of Harvest |
| --- | --- | --- | --- | --- |
| **Trauma Donors** |  |  |  |  |
| A | 84 | Female | Trauma – Hip hemiarthroplasty | Proximal femur |
| C | 46 | Male | Trauma – Pelvic fracture fixation | Pelvis |
| F | 66 | Female | Trauma – Total hip replacement for trauma | Proximal femur |
| H | 75 | Male | Trauma – Pelvic fracture fixation | Pelvis |
| I | 30 | Male | Trauma – Tibial shaft fracture fixation | Proximal tibia |
|  | **Mean Age – 60.2** |  |  |  |
| **Osteoarthritis Donors** |  |  |  |  |
| B | 48 | Female | Total hip replacement | Proximal femur |
| D | 81 | Female |  |  |
| E | 29 | Female |  |  |
| G | 90 | Female |  |  |
| J | 64 | Male |  |  |
|  | **Mean Age – 54.8** |  |  |  |

